# Sex specific molecular responses of Quick-To-Court in Indian malarial vector *Anopheles culicifacies*: conflict of mating and/or blood feeding?

**DOI:** 10.1101/128215

**Authors:** Tanwee Das De, Punita Sharma, Charu Rawal, Seena Kumari, Sanjay Tavetiya, Jyoti Yadav, Yasha Hasija, Rajnikant Dixit

## Abstract

Understanding the molecular basis of mosquito behavioral complexity is central to the design of novel molecular tool to fight against their vector borne diseases. Although, olfactory system play important role to guide and manage many behavioral co-ordinates including feeding, mating, breeding etc., but the sex specific regulation of olfactory responses remains unanswered. From our ongoing transcriptomic data annotation of blood fed adult female olfactory tissue of *A. culicifacies* mosquito, we identified a 383 bp long unique transcript encoding *Drosophila* homolog of Quick-To-Court protein, previously shown to regulate the courtship behavior in adult male *Drosophila*. A comprehensive *in silico* analysis predicts *Ac-qtc* is 1536 bp long single copy gene encoding 511 AA long protein, having high degree of conservation with other insect homolog. Age dependent increased expression of putative *Ac-qtc* in the naïve mosquitoes correlates the maturation of olfactory system, necessary to meet sex specific conflicting demand of mating (mate finding) vs. host-seeking behavioral responses. Though, 16-18 hour of starvation did not altered *Ac-qtc* expression in both the sexes, however blood feeding significantly modulated its response in the adult female mosquitoes, confirming that it may not be involved in sugar feeding associated behavioural regulation. Finally, a behavioural-cum-molecular assay indicated that natural dysregulation of *Ac-qtc* in late evening may promotes key mating event of successful insemination process. We hypothesize that *Ac-qtc* may play unique role to meet and manage the sex specific conflicting demand of mosquito courtship behaviour and/or blood feeding behaviour in the adult female mosquitoes. A molecular mechanism elucidation may provide new knowledge to consider *Ac-qtc* as a key molecular target for mosquito borne disease management.

## Introduction

Mosquitoes are vectors for many deadliest infectious diseases including Malaria, Dengue, Chickungunya, Zika Virus infection, Yellow fever etc. Pathogen transmission is caused by adult female mosquito when it sucks blood from a suitable vertebrate host. Thus suppression of mosquito population and interruption of the mosquito-human interaction by means of chemical insecticides still play vital role to control vector borne diseases. However, fast emergence of insecticide resistance possesses a challenge to design alternative molecular tools to interfere complex feeding and mating behavioral properties [1]. Compared to the vast research on female mosquitoes, male mosquito biology is poorly explored possibly due to its indirect influence on parasite transmission. But males contribute significant effect on disease transmission by inducing several behavioural changes in females and thus maintaining mosquito life cycle continuity. In nature both male and female feed on nectar sugar for their regular metabolic energy source; however the molecular basis of evolution and regulation of dual feeding behavior i.e. nectar sugar vs. blood meal in adult female mosquitoes is not fully understood [2-3]. Likewise, clarifying that how mosquitoes manages complex mating behavioral events i.e. swarm formation, suitable mate finding and successful aerial coupling is yet a major challenge to entomologist [4-7]. Though, in *Drosophila melanogaster* a few molecular markers linked to mating behavior have been characterized [8-10].

Neuro-olfactory system regulate many complex behavioural responses including feeding, mating, host seeking and blood feeding in the mosquitoes [11-14]. Though, current evidence indicate that both male and female mosquitoes olfactory system encodes fairly similar number of proteins [15-16], but it is not known that how sex specific olfactory proteins manage conflicting demand of feeding and/or mating behavior. A modest changes occurs in the olfactory repertoire in response to blood feeding in adult female mosquitoes [17-18]. Owing to these observation it is plausible to hypothesize that sex specific regulation of some of the unique genes may facilitate and manage similar functions e.g. mate partner location by adult male/female mosquitoes or host finding for blood feeding by adult female mosquitoes.

*A. culicifacies* is one of the major rural vector in India, responsible for >65% of malaria cases and controlling this mosquito species became worse due to rapid emergence to multiple insecticide resistance [19]. While understanding the complex feeding behavior of adult *A. culicifacies* female mosquito [3], unexpectedly we identified a unique transcript encoding ‘*quick to court’* (QTC) protein, a homologue of Drosophila QTC (Q9VMU5), from the olfactory system of blood fed mosquito (Unpublished). The *quick-to-court* gene encodes a predicted coiled coil protein which play important role to drive the male courtship behavior in *D. melanogaster,* and predominantly express in olfactory organs, central nervous system and male reproductive tract [20-21]. Mutations in the *Dm-qtc* not only cause males to show elevated levels of male-male courtship, but also favored abnormally quick courtship, when placed in the presence of a virgin female [18]. Recently, a QTC homolog has also been identified from whole body transcriptome of the insect *Bactrocera dorsalis* [22-23] but their function remains uncharacterized so far.

A significant modulation (∼5 fold up regulation) in response to blood feeding in the olfactory tissue prompted us to understand the possible role of this unique transcript named *Ac-qtc* in managing sex specific conflicting demands of ‘mate choice’ and/or ‘food choice’ in the mosquito *A. culicifacies*. A comprehensive *in silico* function prediction analysis and extensive transcriptional profiling establish a possible correlation that *Ac-qtc* may play a crucial role in the regulation and co-ordination of mosquito sex specific differential behaviors.

## Materials & Methods

Fig. 1a represents a technical overview and work flow of the experiments to demonstrate the sex specific possible role of *Ac-qtc* in adult mosquito *A. culicifacies*.

**Figure 1:**
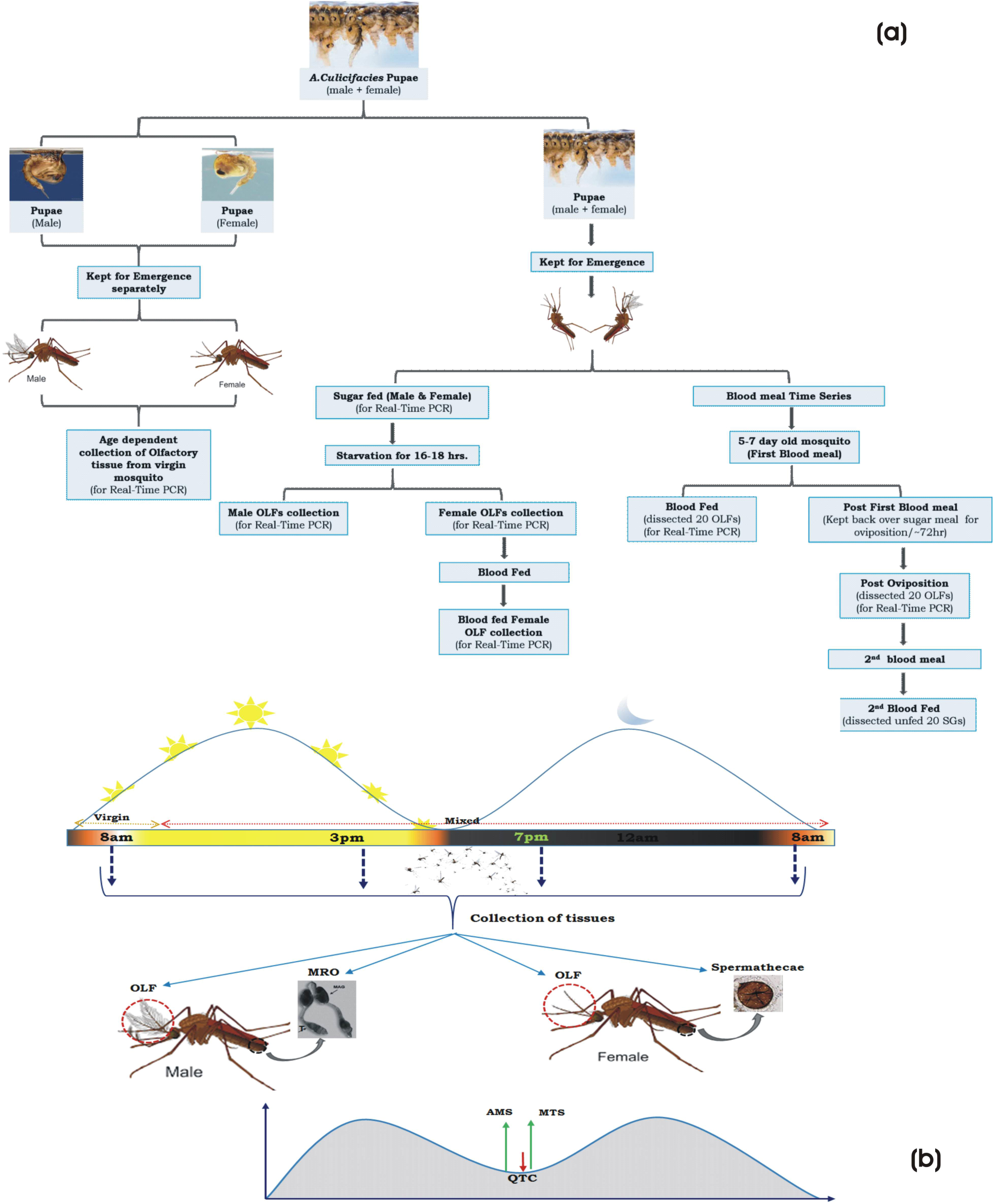
Technical designing and work flow of the experiments. (a) Experimental overview to demonstrate the possible role of *Ac-qtc* in mosquito behavioural regulation. (b) Pictorial presentation of the assay that was performed to correlate the function of *Ac-qtc* in the mating success of *A.culicifacies* mosquito.

### Mosquito Rearing and Maintenance

Cyclic colony of the mosquito *A. culicifacies*, sibling species A are reared and maintained at 28±2°C, RH=80% in the central insectary facility as mentioned previously [3, 24]. All protocols for rearing and maintenance of the mosquito culture were approved by ethical committee of the institute.

### Tissue collection and RNA extraction

From the ice anesthetized adult *A. culicifacies* mosquitoes, the desired tissues viz. olfactory (which include antennae, maxillary palp, proboscis and labium), brain, reproductive tissues (which include testes and male accessary gland for male reproductive organ, and spermathecae/autrium for female reproductive organ), legs of both male and female mosquito were dissected and collected in Trizol. Total RNA were isolated by the standard Trizol method as described previously [3, 25].

### Bioinformatics Analysis

The putative *Ac-qtc* was identified as a partial cDNA from olfactory tissue cDNA library sequence database of the blood fed adult female mosquito (Unpublished). Multiple BLAST analysis against mosquito draft genome and other transcript database was done using analysis tools available at www.vectorbase.org. Domain prediction, multiple sequence alignment and phylogenetic analysis was done using multiple software as described earlier [25].

### Behavioural-cum-molecular assay

To track the possible role of *Ac-qtc* in mating behavior, we designed an assay (Fig.1b) favouring sex specific changes of the behavioural activities occurring in response to day/night cycle in the 5-6 days old mosquitoes. As per assay design we collected olfactory and reproductive tissues from either virgin and/or mixed cage mosquitoes of both the sexes. For mating success experiment, first we mixed at least 100 male/female virgin mosquitoes in a single cage at early morning (8:00am) and collected tissues at 3:00pm, 7:00pm and overnight/8:00am next morning from the same cage. To test and validate the insemination process completion we profiled and compared the expression of two independent sperm specific transcripts in the spermethecae of adult female mosquitoes. Previously characterized sperm specific genes (AMS/FJ869235.1and MTS/FJ869236.1) of the mosquito *A. gambiae* **[26]** were queried to search and select sperm specific homologue from the draft genome database of the mosquito *A. culicifacies*. The identified *Ac-ams* (ACUA010089) and *Ac-mts* (ACUA014389) transcripts sequences were used to design RT-PCR primers.

### cDNA Preparation and Gene Expression Analysis

1 µg of Total RNA were used for first strand cDNA synthesis using Verso cDNA synthesis Kit (Thermo Scientific) by following the standard manufacturer protocol. Routine differential gene expression analysis were performed by the normal RT-PCR and agarose gel electrophoresis protocol. For relative gene expression analysis, SYBR green qPCR (Thermo Scientific) master mix and Illumina Eco Real-Time PCR machine were used. PCR cycle parameters involved an initial denaturation at 95^o^C for 5 min, 40 cycles of 10 s at 95^°^C, 15 s at 52^°^C, and 22 s at 72^°^C. Fluorescence readings were taken at 72^°^C after each cycle. The final steps of PCR at 95^°^C for 15 Sec followed by 55^°^C for 15 sec and again 95^°^C for 15 sec was completed before deriving a melting curve. To better evaluate the relative expression, each experiment was performed in three independent biological replicates. Actin gene was used as internal control in all the experiment and the relative quantification data were analyzed by 2^−ΔΔCt^ method [27]. For statistical analysis of differential gene expression student’s test was used.

## Results and Discussion

### Identification, annotation and molecular characterization of *Ac-qtc*

To identify the differentially expressed genes in naïve sugar fed *vs.* blood fed olfactory system (OLF) of *A. culicifacies* mosquito, currently we are annotating large scale RNA-Seq transcriptomic databases (Unpublished). Unexpectedly we identified a unique transcript from blood fed olfactory transcriptome encoding Drosophila homolog of Quick-To-Court (QTC) protein, which play crucial role in many aspects of male mating behavior. BLASTX analysis of the partial 383 bp long transcript showed 59% identity with the *qtc* homolog of *Drosophila melanogaster*, but no putative conserved domain was identified. Thus to retrieve full length *A. culicifacies qtc* transcript we performed BLASTN analysis against *A. culicifacies* genome and transcript databases using partial transcript as a query sequence. Comparative alignment analysis of the partial and full length transcript indicated that the putative *Ac-qtc* transcript lacks both 5’ and 3’ sequences. Therefore, for detail *in silico* analysis we proceed with the full length *qtc* transcript (ACUA027268) having 1536 bp nucleotide which encodes a 511 amino acid long QTC protein, carrying the coiled-coil domain signature at the 3’ end of the sequence. Quick to court is a single copy gene, comprised of 50 bp 5’ UTR region followed by five exons and four introns followed by 50 bp 3’UTR region [Fig.2a]. A comprehensive primary structural analysis of this full length transcripts (ACUA027268) revealed that it has four coiled-coils features and one conserved GRIP domain at the 3’ end of the sequence [Fig.2b; Supplementary table 1]. Multiple sequence alignment (Fig. S1) and phylogenetic analysis revealed a high degree of sequence conservation within the mosquito and other insect species [Fig.2c]. RT-PCR analysis indicated *Ac-qtc* constitutively express in all the aquatic stages of development, except egg of the mosquitoes [Fig.2d], suggesting that this quick-to-court protein is evolutionary conserved, with common function in the regulation of insects behavioural responses.

**Figure 2:**
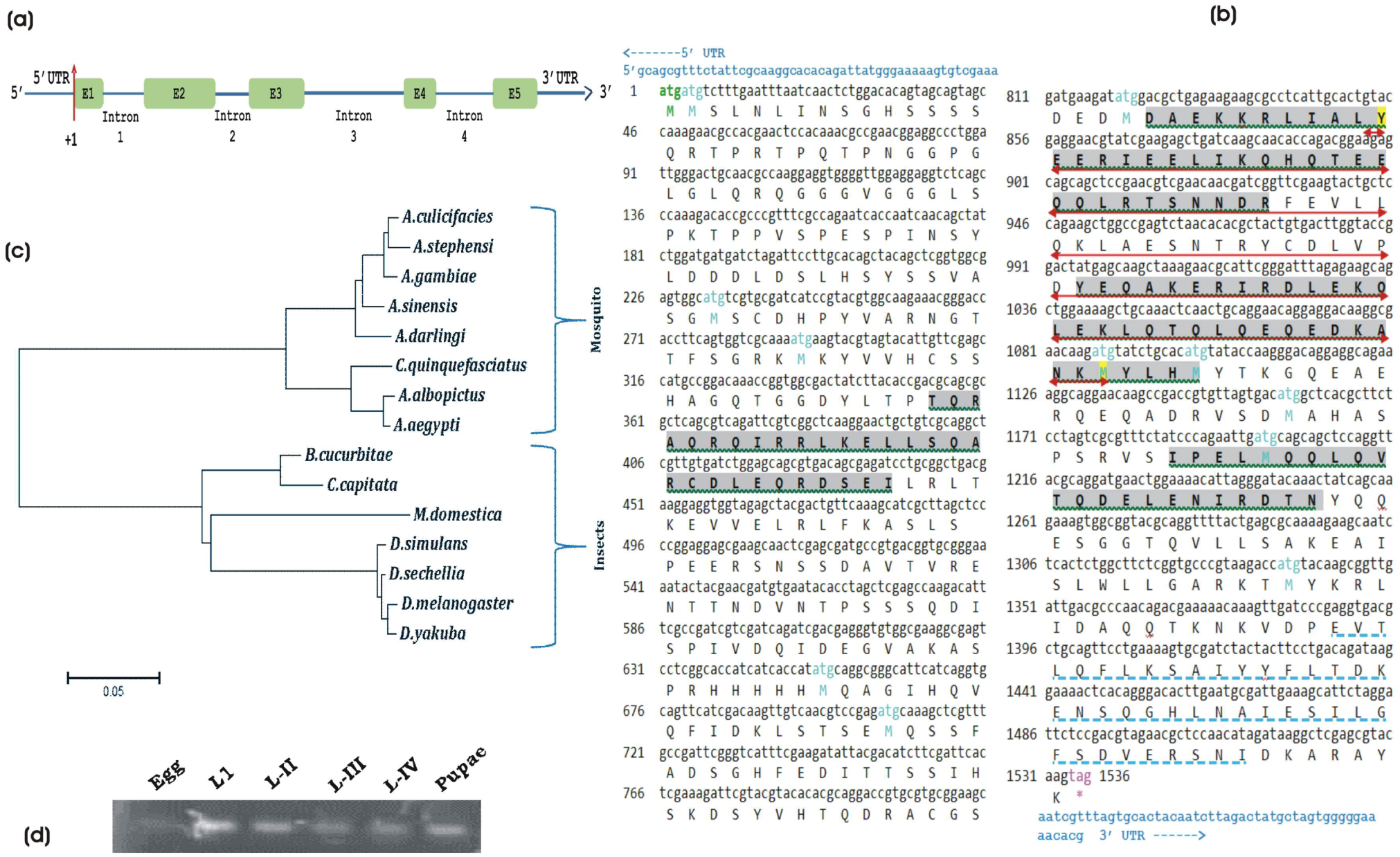
Genomic and molecular Gene Organization of *A. culicifacies quick-to-court* gene:-(a) Schematic representation of the genomic architecture of the mosquito *Ac-qtc.* Five green colored boxes indicates the introns and +1 mark the transcription initiation site. (b) Gene organization and molecular features of full length *Ac-qtc*: The gene contains 1536 bp nucleotide, encoding 511 AA long peptides with four coiled-coils domains. Both 5’ and 3’-UTR regions are highlighted (Yellow) and shown in bold & capital letters. The complete coding region of 511 amino acids starts from ATG/Methionine/green color, ending with TAG/Red/*. The different features are highlighted with different color code, viz. Coiled-coils domain (grey color and green underlined), Pre-folding domain (Red arrow) and GRIP-domain (sky blue dotted line). (c) Phylogenetic relationship of *Ac-qtc* with other mosquito species and Drosophila indicates that *A. culicifacies qtc* is clustered within the mosquito domain and have much greater similarity with mosquito *qtc* than Drosophila homologs. (d) RT-PCR mediated developmental expression of *Ac-qtc* in *A. culicifacies.*

### *Ac-qtc* abundantly express in the olfactory and brain tissue of adult mosquitoes

The molecular characterization of *qtc* gene is limited to the model insects *D. melanogaster* only, where it expresses in the olfactory organs, central nervous system of both the sexes and male reproductive tract, possibly to regulate male courtship behavior [18]. However, so far it remains uncharacterized from any mosquito species. Sex and tissue specific transcriptional profiling of *Ac-qtc* in naive mosquitoes revealed that male mosquito has dominant expression of *qtc* gene in all of the tissue including olfactory tissue, brain, reproductive organs etc. as compared to the female [Fig 3]. Interestingly, olfactory tissue of male *A. culicifacies* mosquito showed relatively higher abundance than other tissue indicating its possible involvement in the regulation of mosquito behavior.

**Figure 3:**
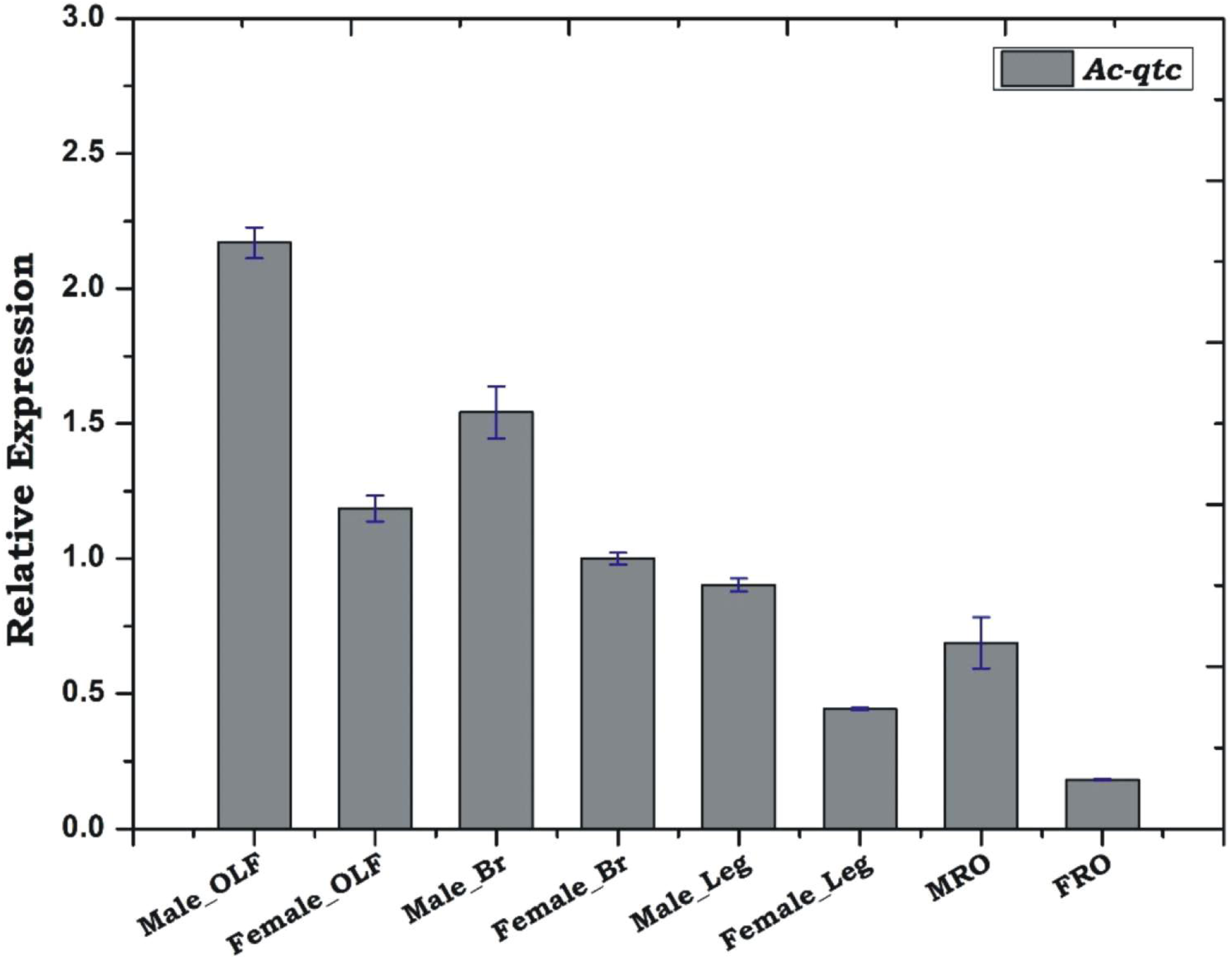
Tissue specific transcriptional behaviour of *Ac-qtc*. Tissue specific relative expression analysis of *Ac-qtc*. Male OLF: Male olfactory tissue (Antennae, maxillary palp and proboscis); Female OLF: female olfactory tissue; Male-Br: Male brain; Female-Br: female brain; Male-leg; Female-leg; MRO: Male reproductive organ; FRO: Female reproductive organ.

### Sex specific age dependent expression may regulate olfactory system maturation

Unlike *Drosophila*, unavailability of the proper molecular marker restricts our understanding of the complex mating biology in mosquitoes. In case of the mosquito *A. culicifacies* the behavioural studies on mating behaviour are too limited [28]. However, our unexpected finding of the *quick-to-court* transcript from the blood fed olfactory tissue transcriptome data allowed us to hypothesize that *Ac-qtc* may be one of the key molecular factor to derive sex specific behavioural modulation in the mosquito *A. culicifacies.*

To test this hypothesis, first we examined sex specific relative expression of a pool of four transcripts including *qtc*, newly identified from the ongoing olfactory tissue transcriptomic study of blood fed adult female mosquito (Fig. S2), revealing that male olfactory system mature faster than female mosquito of same age. Next, we monitored age dependent transcriptional regulation of *Ac-qtc* in virgin male and female mosquitoes. It showed age dependent enrichment of *Ac-qtc* at a highest level (∼6 fold up regulated/ *p*≤ *0.004*) on 5^th^ day in both sexes, when compared with 1 day old virgin mosquitoes [Fig. 4a, b], followed by at least 2 fold down regulation (p≤ *0.0172*) on 7^th^ day. Interestingly post 7^th^ day *Ac-qtc* expression switched to up regulation in male mosquitoes, but remains constant after 5^th^ day in case of female mosquitoes. A similar pattern of *Ac-qtc* gene expression could also be observed in the possibly mated male mosquitoes [Fig. 4c], which were collected from mixed cage containing equal number of male and female mosquitoes.

**Figure 4:**
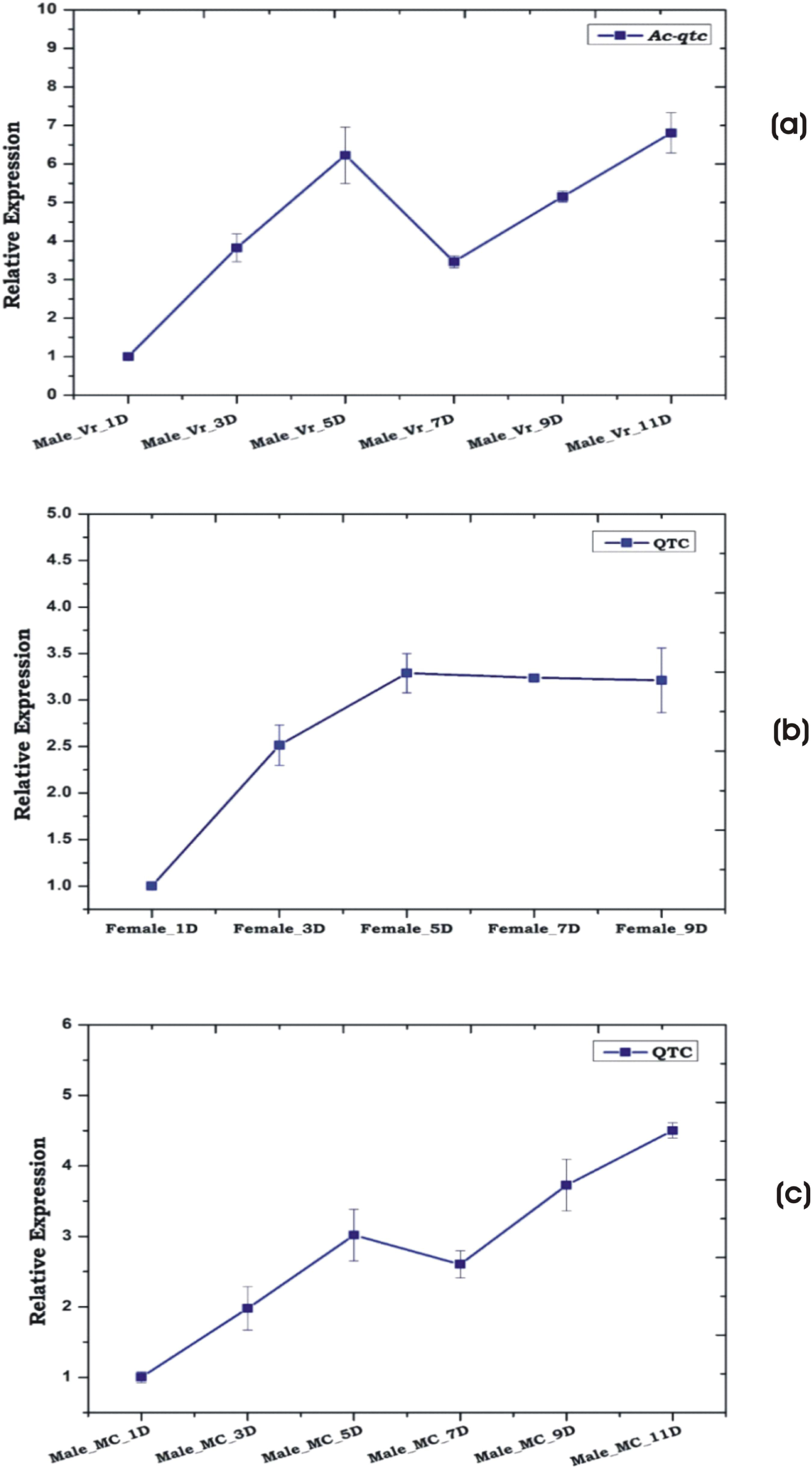
Age dependent transcriptional profiling of *Ac-qtc* in male and female mosquito. (a) Age specific relative transcriptional regulation of *Ac-qtc* in virgin male mosquito. Male-Vr-1D: Male virgin mosquito of 1Day old. (b) Transcriptional regulation of *Ac-qtc* in virgin Female mosquito according to their age. Female-Vr-1D: Female virgin mosquito of 1Day old. (c) Age dependent relative expression analysis of Ac-qtc transcript in the possibly mated mosquito. Male-MC-1D: Male mosquito collected from mixed cage (MC) containing equal number of male female mosquito.

Although, it is yet un-clarified that how environmental guided non-genetic and/or genetic factors regulate the complex sexual behavioral events, but above observations indicated that once males achieved specific age of adulteration, the dys-regulation of *Ac-qtc* by unknown mechanism may promote the courtship behaviour (see next para). These results also corroborate with the previous findings in *Drosophila* where mutation in *Dm-qtc* gene causes accelerated male-male courtship behavior [18]. Recent study by *Houot et. al.* also suggest that *qtc* and *shaker* genes, which abundantly expresses in the neuro-olfactory system of the *Drosophila*, causes decreased ability to discriminate between the sex targets/ probably due to the decline perception to wild type female pheromone [28-31].

### Natural dysregulation of *Ac-qtc* may promote mating success

Alternatively to the above observations we also interpreted that a significant down regulation (p≤ *0.0172*) of *Ac-qtc* in the 5-7 day old virgin adult male mosquitoes, may be crucial for the auto-activation of courtship behavior in case of *A. culicifacies* mosquitoes. In the lack of any molecular marker for mating behaviour studies, we attempted to trace the possible functional correlation of male *Ac-qtc* in the mating success i.e. insemination event completion in the laboratory reared mosquitoes. However, our initial experiments of *Ac-qtc* gene silencing using purified dsRNA injection in the thorax of male mosquito’s remains unsuccessful, primarily due to high mortality rate of these mosquitoes (data not shown). Available literature suggests that most anopheline mosquitoes mating behavioural activities commenced at the onset of sunset, usually at 17:00 pm which may continue till 20:00 pm [29, 32-33]. We hypothesize that if true, the transcriptional modulation of *Ac-qtc* in response to dawn/dusk cycle, must have functional correlation with the mating success, especially insemination events where adult females receive and store the sperms in her spermethecae delivered by male during copulation [34]. To test this idea, first we identified two sperm specific transcripts from the draft genome of the mosquito *A. culicifacies*, using (*ams* & *mts*) as query sequences, previously characterized from *A. gambiae* [26]. To validate sperm specificity, the primers designed against *Ac-ams* (ACUA010089) and *Ac-mts*/ACUA014389) were tested by RT-PCR in the male accessory glands (MAG) and spermathecae (SPT) collected from laboratory reared 3-4 day old virgin male and female mosquitoes respectively (Fig 5a). A non-specific poor amplification visible in case of *Ac-ams* in virgin female mosquito spermethecae was verified by incorrect melting curve signal appeared in Real-Time PCR data (Supplemental Fig. S3). Next, to trace the possible functional correlation of *Ac-qtc* in mating success, we analysed and compared sex specific regulation of *Ac-qtc* in the olfactory tissue, and sperm specific *Ac-ams/Ac-mts* genes in the male reproductive organs (including MAG and testis) and spermathecae (SPT) tissues, collected two hours prior or later onset of the sunset as described in methodology section (See experimental design Fig. 1b).

**Figure 5:**
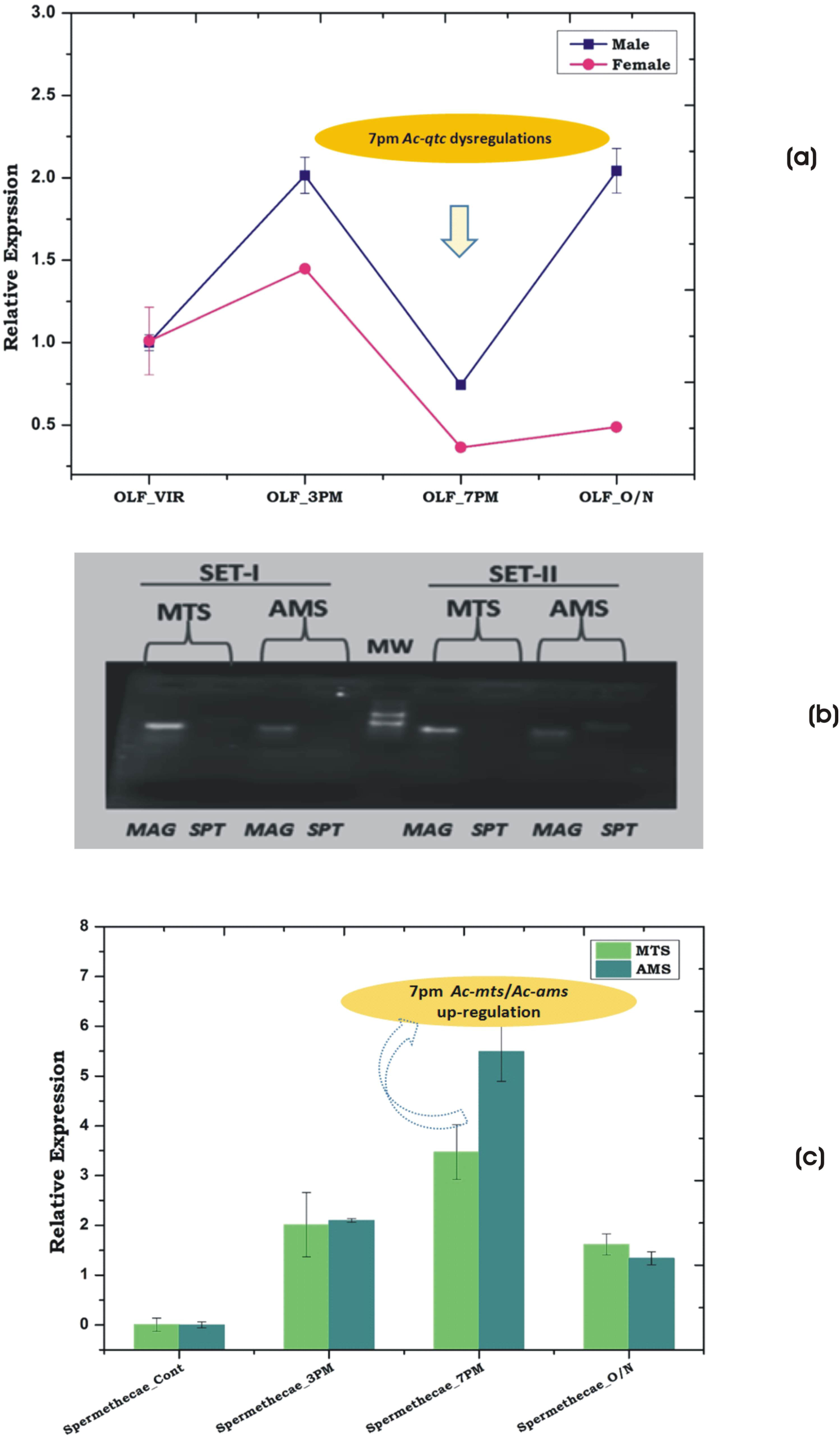
(a) RT-PCR mediated expression profiling of *Ac-ams* and *Ac-mts* in MAG and Spermethecae. MAG (Male Accessory Gland). (b) Transcriptional regulation of *Ac-qtc* at different circadian time in the olfactory tissues of both male and female *A. culicifacies* mosquitoes. (c) Transcriptional response of *Ac-mts* and *Ac-ams* in the spermethecae of *A. culicifacies* at different circadian time.

When compared to the virgin counterpart, significant down-regulation of *Ac-qtc* in the olfactory system in both the sexes at 7:00 pm **(Fig. 5b)** indicated that lower expression *Ac-qtc* may favour the increased mating frequency and active courtship engagement. Though molecular mechanism is yet to be understood, however, interestingly significant modulation of *Ac-ams/Ac-mts* expression in the mated female spermethecae **(Fig. 5c)** as well as male reproductive organs (Fig. S4) supports the idea that a natural dysregulation of *Ac-qtc* in the late evening i.e. 7:00 pm may favours the increased mating frequency facilitating the release of unknown sex driving factors for successful insemination event completion in the copulating couples.

### *Ac-qtc* regulation is independent to nutritional status of the mosquitoes

Proper feeding and mating are two unique but interdependent behavioral properties of any biological system that facilitate their survival and reproductive success synergistically. The molecular basis of the sex specific regulation of these conflicting behavioral demands remains largely unknown. Current studies in *Drosophila* suggest that food odor and sex specific pheromone signals in the neuro-olfactory system may facilitate to drive the meal and/or mate attraction [31]. Thus to test whether *Ac-qtc* have any sex specific relation to the nutritional status of naïve adult mosquitoes, we examined and compared the relative expression of *qtc* transcript in starved vs. sugar fed mosquito in both the sexes. To perform this experiment, we collected olfactory tissue either from 5-6 day old sugar fed or 16-18 hrs starved mosquitoes. Our relative gene expression analysis indicated that starvation does not affect the abundance of *qtc* transcript, but showed a twofold up-regulation (p≤ *0.03*) in response to immediate i.e. 1-2 post blood feeding [Fig. 6]. Together these data, suggested that *Ac-qtc* gene may not be essential for regulating mosquito sugar feeding behavior but may have play important role in the regulation of host seeking/blood feeding behavior in the adult female mosquitoes.

**Figure 6:**
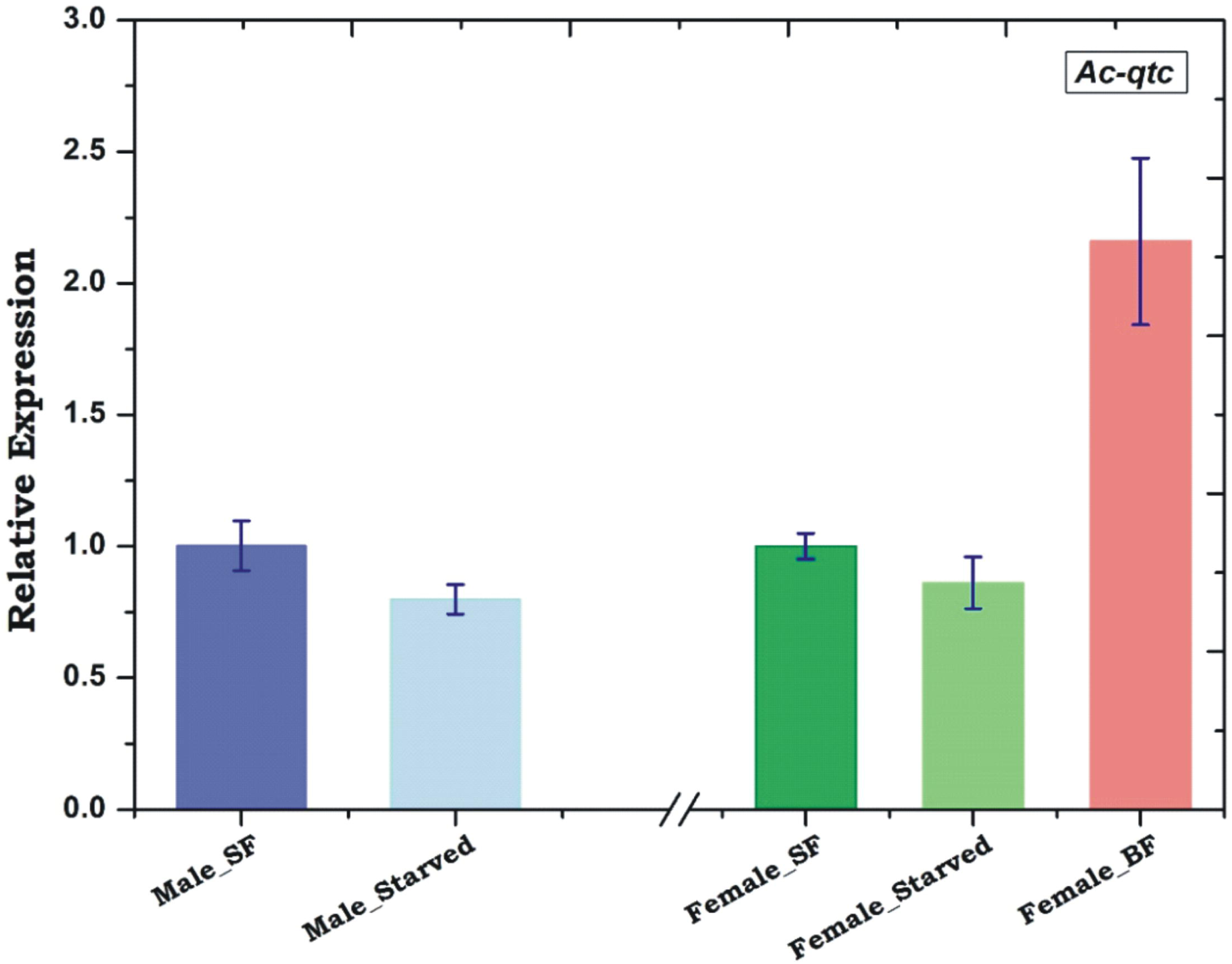
Effect of starvation on *Ac-qtc* expression. Both male and female mosquitoes were kept starved for 16-18 hr and the olfactory tissue were collected for *qtc* expression study. Additionally blood meal was provided to female mosquitoes after starvation.

### Blood meal alters expression of *Ac-qtc* in adult female mosquito

Originally, we identified the unique transcripts *Ac-qtc* from 30hr post blood fed adult female mosquito RNAseq library, and above experimental validation (Fig. 6) prompted to further evaluate *Ac-qtc* role in response to blood feeding behavior of female mosquitoes. To clarify this complexity, we performed a blood meal time series experiment as described earlier [3], in which we collected olfactory tissue of *A. culicifacies* mosquito depending on their age and blood feeding status. *Ac-qtc* expression analysis revealed increased abundance till 5^th^ day, but significantly (2.5 fold / p≤ *0.03*) down regulated on 7^th^ day in the naive unfed adult female mosquitoes, a pattern similar to the adult male mosquitoes (Fig. 5), which may probably be to attend and meet an optimal courtship behavioral success in both the sexes.

Although unclear whether first blood meal transiently and/or completely pauses re-mating events, however, interestingly, a consistent up regulation of *Ac-qtc* just after blood feeding (within 30 min) till 30hrs post blood meal (Fig. 7), indirectly suggest that at least for first 30 hrs post blood meal, adult female mosquito may not seek any courtship event. Furthermore, a continuous sharp down regulation (∼4 fold) of *Ac-qtc* till 72 hrs post first blood meal, remains a questionable observation that whether *Ac-qtc* promotes host-seeking behaviour for second blood meal and/or mate partner finding success, because after oviposition and prior second blood meal we noticed significant up regulation (∼1.3 fold; Fig. 7). Second blood meal again rapidly down regulated the *Ac-qtc* expression, when examined 30hr post blood meal in the olfactory tissue, together indicating that *Ac-qtc* may have unique role in deriving dual mode of behavioural responses possibly to meet the conflicting demand of sexual mate partner and/or suitable vertebrate host finding for blood feeding.

**Figure 7:**
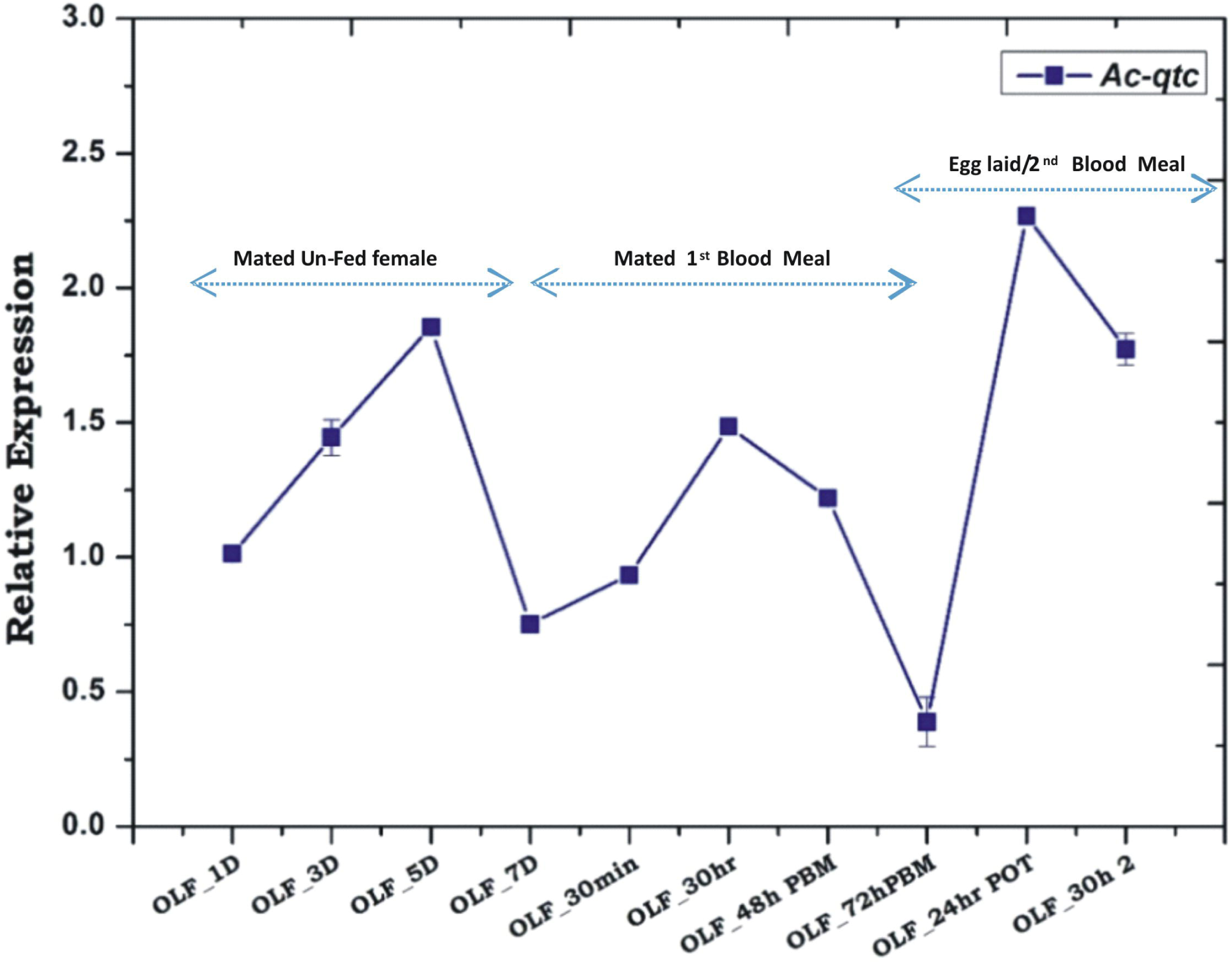
Transcriptional behavior of *Ac-qtc* in two consecutive blood meal: Age dependent blood meal responses of *Ac-qtc* transcript in the olfactory tissue of adult female mosquito *A. culicifacies.* OLF-1D – OLF-7D: Olfactory tissue from 1Day -7 Day old female; OLF-30min: Olfactory tissue collected from 30min post blood fed mosquito; OLF-30hr: Olfactory tissue collected from 30hr post blood fed mosquito; OLF-48hr: Olfactory tissue collected from 48hr of post blood fed mosquito; OLF-72hr: Olfactory tissue collected from 72hr post blood fed mosquito; OLF-24hr POT: Olfactory tissue collected from 24hr of post oviposition of mosquito; OLF-30hr 2: Olfactory tissue collected from 30hr of 2^nd^ blood meal.

## Conclusion

Understanding the sex specific molecular genetics of mosquito’s behavioural biology is more complex. This is partly due to unique nature of blood feeding evolution and adaptation in the adult female mosquitoes, and partly due to limitation of generating mutants. Through comprehensive molecular approach, we examined the sex specific transcriptional regulation of a unique transcript *Ac-qtc*, and predicted its possible role with ‘food choice’ and/or ‘mate choice’ behavioural performance **(Fig. 8).** Our data provide the first molecular evidence that Ac-QTC proteins may have dual mode of action in the regulation of cluster of mosquito olfactory genes that are linked to mating success and/or blood feeding in adult female mosquitoes. We believe these findings may guide to uncover the functional nature of *Ac-qtc* controlling complex mating and/or blood feeding behaviour in mosquitoes.

**Figure 8:**
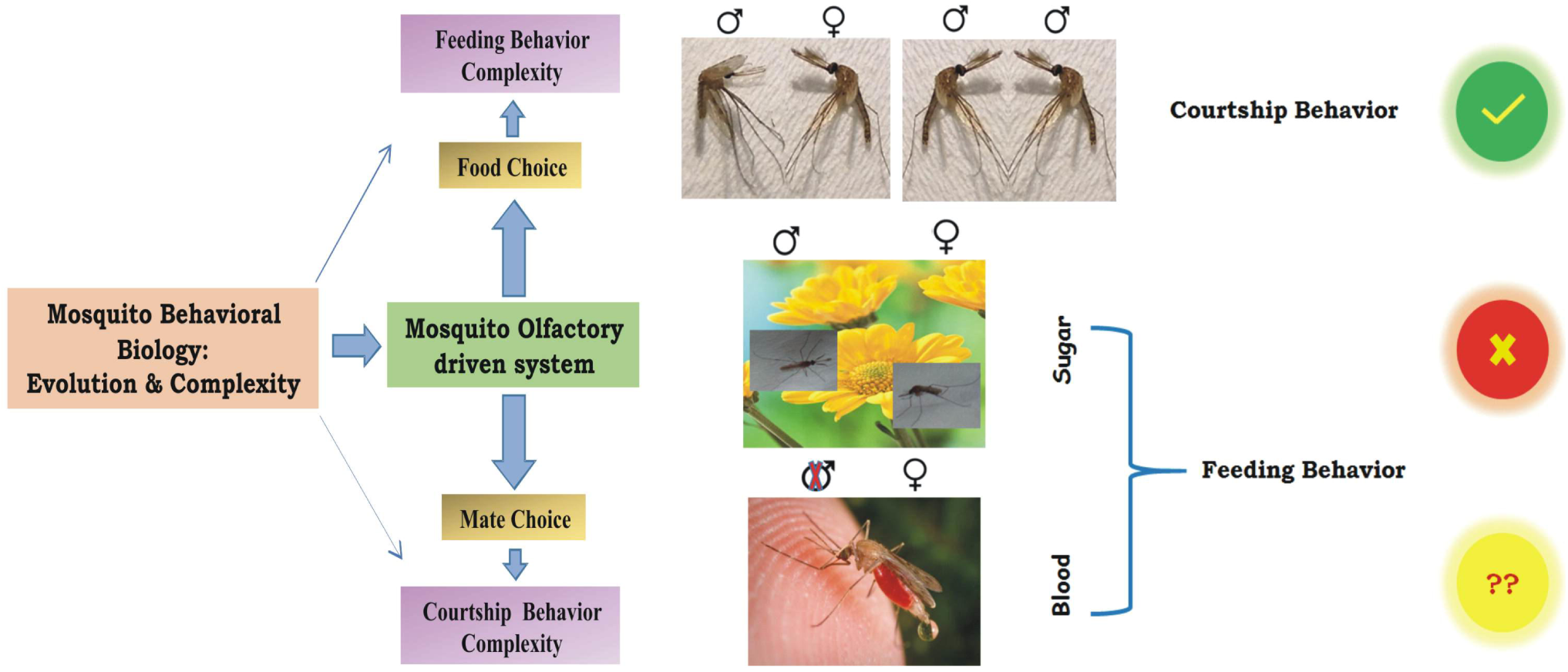
Proposed hypothesis for the possible function of quick-to-court gene in mosquito.

### Sequence data submission

Identified partial CDS sequence of *Ac-qtc* submitted to NCBI Genbank with Accession no. KX575650.

## Declarations

### Authors contribution statement

Conceived and designed the experiments: TDD,RD; Performed the Experiments: TTD, PS, CR, SK, ST, JY;

Analyzed the data: TDD, YH, RD; Contributed reagents/materials/analysis tools: YH, RD; Wrote the paper: TDD, RD

## Acknowledgement

We thank insectary staff members for mosquito rearing. We also thank Kunwarjeet Singh for technical assistance in laboratory. Finally, we are thankful to Xceleris Genomics, Ahmedabad, India for generating NGS sequencing data.

